# Response to Kozlov *et al.*: Inaccurate estimation of biases in herbarium specimen data

**DOI:** 10.1101/2020.09.01.278606

**Authors:** Emily K. Meineke, Charles C. Davis, T. Jonathan Davies

## Abstract

Kozlov and colleagues^1^ call into question the application of herbarium specimens to quantify historical patterns of herbivory^2–5^. It is already widely appreciated that collectors of herbarium specimens may tend to avoid insect damage, thus making herbivory estimates from herbarium specimens potentially down-biased^2^. However, Kozlov *et al.* additionally suggest that variation in sampling selectivity among collectors and curators may lead herbarium specimens to misrepresent patterns of herbivory in nature. The authors sought to quantify these biases by collecting and contrasting insect herbivory data across 17 plant species from herbarium versus standard field ecological sampling procedures, and then assessed the selection of these specimens by curators. They concluded that herbivory estimates from herbarium specimens are highly variable, rendering them an inaccurate representation of herbivory in nature. Our re-analysis of Kozlov *et al.*’s data suggests that, in contrast with their results, herbarium specimens indeed provide a useful record of herbivory as long as sample sizes are appropriate. In addition, we assert that by arguing that herbarium specimens are “distorting mirrors”, Kozlov *et al.*’s conclusions fundamentally overstep their data, which narrowly assesses biases across species. Kozlov *et al.* argue that herbarium specimens are inaccurate data sources, but fail to characterize the specific circumstances under which assumed biases would apply. Thus, Kozlov *et al.*’s data do not support their main premise, and the authors extrapolate beyond the specific biases investigated in their study; we believe their contribution does a disservice to researchers interested in exploring the potential value of herbarium specimens for studying herbivory through time.

## Main text

Kozlov *et al.* ^1^ assert that estimates of herbivory damage derived from herbarium specimens are unreliable. Assessing leaf area removed by insects to infer herbivory is a common practice^6^. However, it is well established that insect damage is distributed unevenly; some plants experience extensive damage, whilst others exhibit little or no signs of herbivory (e.g.^7^). It is critical, therefore, to sample multiple individuals when estimating species means, especially when assessing single plant species from a given locality (e.g.^8,9^). Kozlov *et al.* gathered too-few and too-variable specimens (n=1 to n=10 for ecological samples, and n=3 to n=20 for herbarium samples) to infer reliable estimates of herbivory. The pathologies resulting from such under-sampling are evident in Fig. 1 presented by Kozlov *et al.* Herbivory estimates derived from ecological versus herbarium sampling vary widely when inferred from only a few herbarium samples, but converge when sample sizes are higher and comparable between sampling strategies. This trend is observable when species are sorted by total sample size (Fig. 1). For example, herbivory estimates for *Betula pubescens* and *Rubus idaeus,* the only focal species in which sample size is high for both categories, align closely (Fig. 1; Appendix S1).

**Figure 1.**
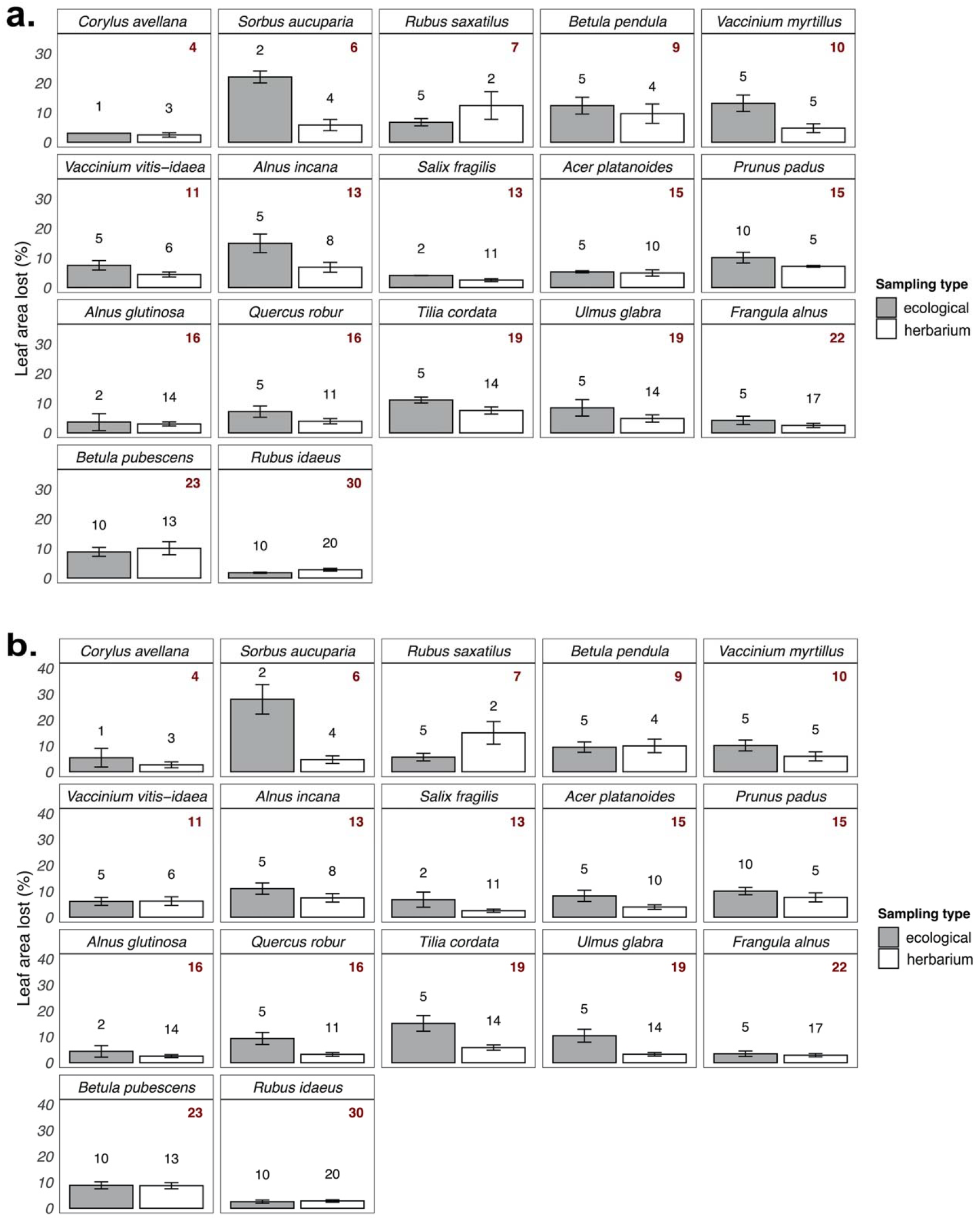
Effects of sample size on herbivory estimates. a) Raw data describing leaf area lost to insect herbivores detected using different sampling methods in Kozlov *et al.*^5,6^. Note that sample sizes differ slightly from those reported in Kozlov *et al.* because of a technical error that caused minor data misreporting in the original paper’s supplement and in the main text figures. b) The same plot with least-squared (marginal) means. This figure includes the exact data from Fig. 1 in Kozlov *et al.*, wherein they report marginal means for which calculation was not included in the original manuscript. For more discussion on why we chose to include panel b), please see Appendix S1.

Using Kozlov *et al.*’s published data, we show that herbivory estimated from ecological and herbarium specimens are highly correlated as long as species have more than 10 data points (Fig. 2) or if we simply weight by sample size (Table 1). By recreating Fig. 3 from Kozlov *et al.*, we demonstrate that herbivory estimates from these two sampling methodologies exhibit a broadly linear relationship which is disrupted by a few plant species represented by small sample sizes (Fig. 2). We re-test the hypothesis that herbivory measurements extracted using the two methodologies are correlated with three sets of models (Table 1). First, we performed weighted linear models using the *lm* function R^10^ with mean herbivory measured on herbarium samples as the independent variable (*x*) and mean herbivory measured using ecological metrics (*y*) as the dependent variable. The data points were weighted by the total number of herbivory samples Kozlov *et al.* collected for a given species (total n= n of herbarium samples + n of ecological samples). Because these weights allow us to account for uneven sample sizes, we restricted our analyses to arithmetic means, not least-squared means (which the authors claim are another way to reduce effects of uneven sampling across species; see Appendix S1). The resulting models— one of which contains a transformed response variable to improve model fit—display broad agreement between measurements of herbivory derived ecological and herbarium sampling procedures (Table 1, Models 1 and 2; *p*= 0.02, r^2^ = 0.27 and *p*<0.01, r^2^ = 0.33). If we restrict Kozlov *et al.*’s data to include only species for which n≥10—still a small sample size for noisy ecological data—the relationship between these arithmetic means calculated from herbarium and ecological sampling is even stronger in a simple linear regression (Table 1, Models 3 and 5; *p*= 0.01, r^2^ = 0.39 and *p*= 0.01, r^2^ = 0.41), and is nearly significant in a simple linear regression focusing on the marginal means Kozlov *et al.* report (Table 1, Models 4 and 6; Fig. 2b: *p*= 0.06, r^2^ = 0.21, and *p*= 0.05, r^2^ = 0.23). Finally, we ran Deming regressions including plant species with n≥10. The Deming modeling procedure is arguably more appropriate for these data than simple linear regression because it allows for errors in both the *x* and the *y* variables (in this case herbivory measured from herbarium and ecological sampling procedures, which both carry error). Both of these regressions were “significant” in that 95% confidence intervals did not overlap zero (Table 1, Models 7 and 8; fits are plotted in Fig S1). We emphasize the plant species not included in Models 3-8 with n<10 are not statistical outliers in terms of herbivory – e.g., points of high leverage that might be identified using Cook’s distance or other outlier detection approaches. Rather, they are instances in which sample size is low and thus herbivory estimates predictably do not match between ecological and herbarium samples because the data are noisy. In summary, the patterns in the data (Figs. 2 and S1) and the statistical models (Table 1) illustrate that herbarium and ecologically derived data are highly correlated when sample sizes are adequate. Therefore, while Kozlov *et al.*’s data provide evidence that estimates of herbivory on herbarium specimens are down-biased, as has been shown previously^2^, differences between species’ estimates are highly consistent when sampling effort is sufficient, contrary to the author’s published conclusion.

**Figure 2.**
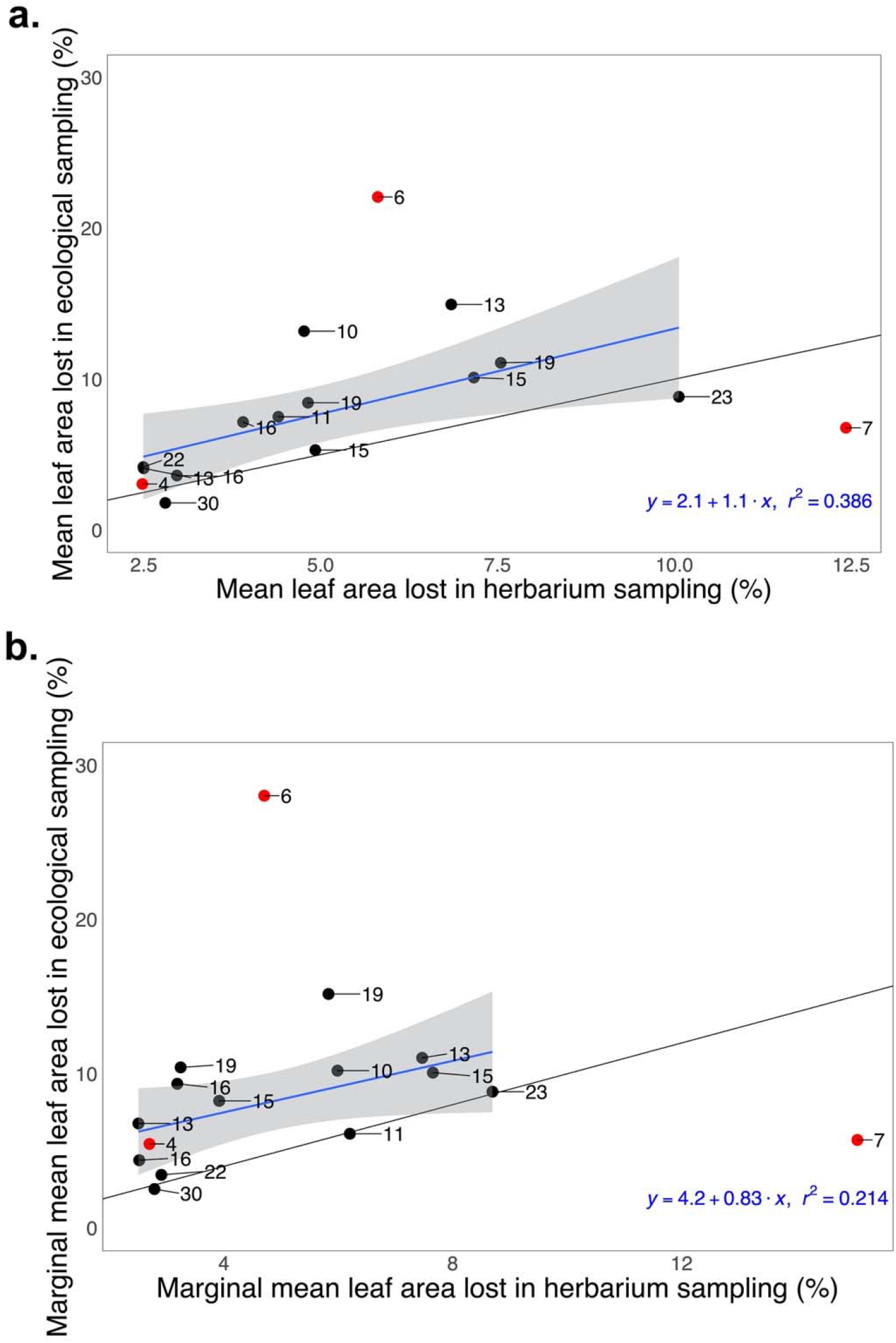
The relationship between herbivory estimated from ecological vs. herbarium specimens. Each data point represents a species, and numbers indicate total sample sizes (ecological + herbarium). For a), the model was derived from the *lm* function in r^10^ and represents the simple linear relationship between mean leaf area loss per species measured in herbarium specimens (*x*) and ecological specimens (*y*). The model fitted to species with ≥10 specimens collected (species in black) is shown in blue (±SE in grey). Associated model parameters are presented in the lower righthand corner (F_1,11_= 8.56, *p*= 0.01). The r^2^ value reported is adjusted r^2^. In b), we show similar patterns in the marginal means calculated by Kozlov et al. for the model that included species for which n>10 (F_1,11_= 4.27, *p*= 0.06). In both plots, the black line represents a 1:1 agreement between herbivory estimates derived from herbarium and ecological samples. Thus, herbarium specimens represent down-biased estimates of herbivory as expected, but these data are nevertheless linearly correlated with herbivory data from ecological sampling of the sample species.

**Table 1.**
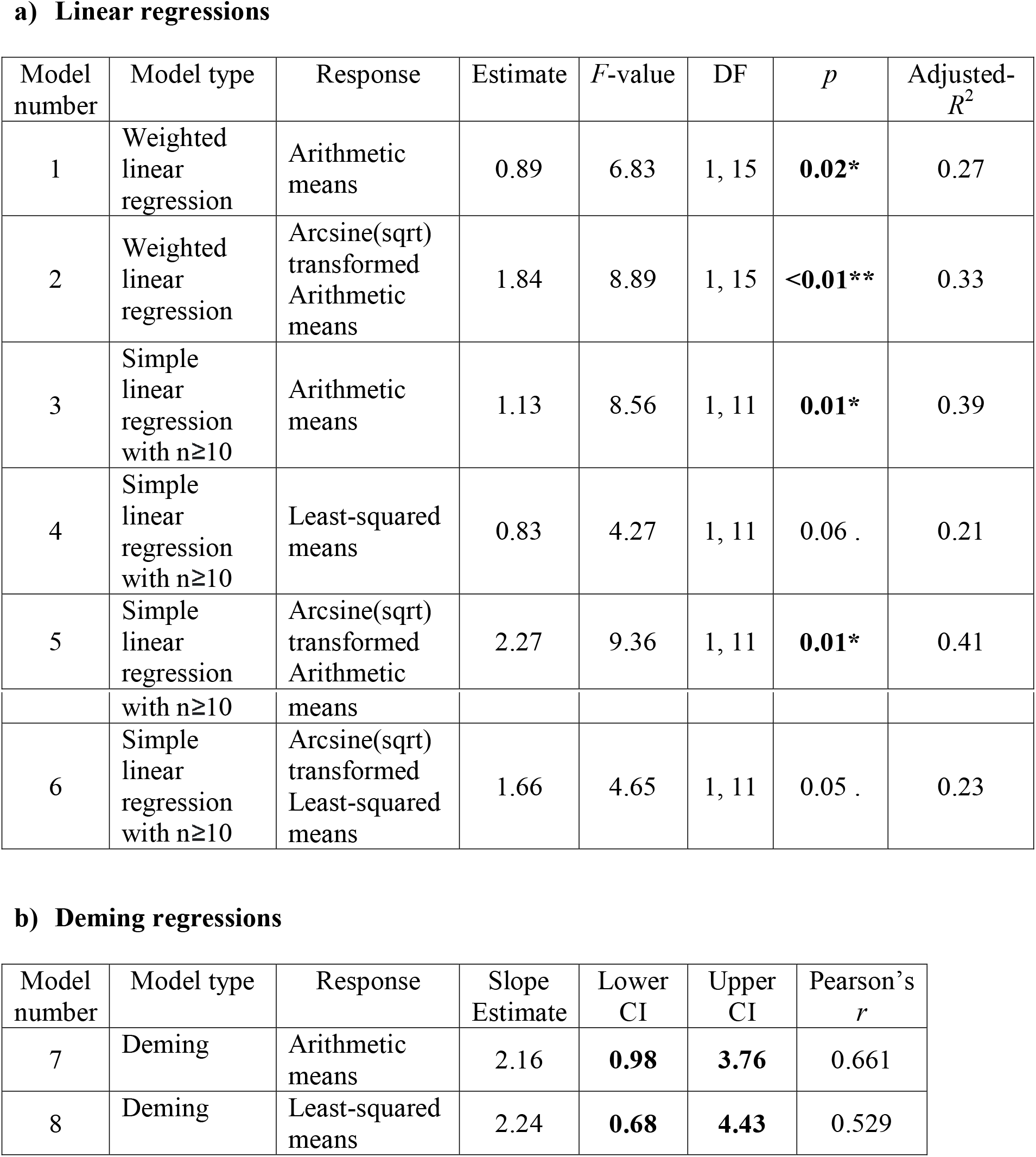
Various models of the relationships between herbivory estimated using herbarium and ecological sampling methodologies. In all models, herbivory estimates derived from herbarium specimens is the independent variable (*x*) and herbivory estimates derived from ecological specimens is the independent variable (*y*). a) As described in the main text, Model 1 has arithmetic means as the response variable, and the data points are weighted by total sample size. Model 2 is identical to Model 1 except that herbivory estimates are treated as proportion data, and the response variable was arcsine(sqrt) transformed to improve model fit. Models 3 and 4 are simple linear regressions, one with arithmetic and one of least-squared mean herbivory estimates as the response variables. Models 5 and 6 are transformed as described above. b) Deming regressions of species with n≥10 data points. Listed values represent 95% confidence intervals. In Deming regression, slope estimates are considered “significant” if the confidence limits do not cross zero, as is the case in our models.

It might be argued that, like Kozlov et al.’s data, herbarium specimens provide few data points if the sampling unit is a given species in a given place at a given time. However, the ecological concept of sample size relates to the independent variable whose effects are being assessed. For example, while just one herbarium specimen from a specific place-species-time combination might be available, if we aim to assess the effects of temperature on herbivory, we can use herbarium specimens from many place-species-time locations with matched temperature data form that location and time period. Further, if the goal is to evaluate the general relationship between temperature on herbivory, we can use hierarchical or mixed effects models that allow us to borrow statistical strength from our better-informed estimates (e.g., species or sites with higher sampling density). Thus, in contrast to most ecological field studies, herbarium specimens offer the opportunity to expand sample sizes vastly across multiple axes, and the effects of individual sites can be captured independently in appropriately fit statistical models that include relevant co-factors. Kozlov *et al.* do not describe any systemic such biases for fellow ecologists to beware of in these herbarium data; however, should they have done so, this information could have been used to usefully inform future model fitting.

Kozlov *et al.* suggest that herbivory estimated from herbarium specimens produces inaccuracies that preclude comparing these metrics among species. However, we know of few studies that use herbarium specimens for this purpose; the focus has instead been on quantifying variability in herbivory over years and space *within* species, and inferring general trends across species. Even if there are idiosyncratic species differences between observed and estimated herbivory from herbarium data, such that herbarium records tend to underestimate herbivory more in some species compared to others, as Kozlov *et al.* suggest, specimens may still serve as an accurate source data on herbivory variation across axes relevant to global change (years, latitude, temperature). For instance, in a comparison of seasonal flowering sensitivity to temperature between observational and herbarium data, Davis *et al.*^11^ demonstrated that while phenological sensitivity measurements from herbarium specimens did not predict phenological sensitivity measured in the field, the effects of seasonal temperature on flowering still aligned. Therefore, differences in the precision with which herbarium data capture herbivory (or phenology) between species does not preclude their use to infer changes in herbivory (or phenology) across environmental gradients or over time. Thus, Kozlov’s criticism, which addresses bias when comparing estimates among plant species, not over time, is not relevant to how most such data are used, including in our study^2^ examining changes in herbivory in the eastern United States over time.

While we have shown that the conclusions drawn by Kozlov *et al.* from the comparison of herbivory estimates on ecological versus herbarium samples are mistaken, better quantifying the true relationships between sampling approaches would be a useful contribution to the literature given the growing number of publications in this area, including some of our own work (e.g.^2,3,12^). In a second analysis, Kozlov *et al.* provide one step toward this goal, polling 17 herbarium curators to assess their biases against the accession of specimens with insect damage. Here they show that herbarium curators are differentially biased against damaged specimens, and thus conclude that herbarium specimens cannot provide reliable estimates of herbivory in nature. We do not dispute that curators likely differ in their selectivity of specimens for vouchering; however, we again disagree the conclusions drawn by Kozlov *et al.*

If curators are inconsistent in their tolerance for damage, and assuming curator bias does not itself vary systematically across the axis of interest, individuals’ preferences would simply add a further source of noise in the data. Although not a source of bias, added noise in herbivory estimates could be problematic in statistical modelling fitting given the natural variability in herbivory damage across individual plants^13^. New machine learning methods provide an exciting path forward by allowing herbivory data to be collected rapidly from greater numbers of specimens^14^, decreasing the noise-to-signal ratio in datasets. We note that, given the inherent differences in collector and curator preferences, it is perhaps more impressive that it is still possible to extract expected patterns in herbivory across space and time from herbarium specimens^3^. However, Kozlov *et al.*’s observation of variability in tolerance to herbivory exhibited by curators does raise an important point regarding differences in cultural practices among herbaria. Though, we are unaware of any studies that explicitly compare herbivory rates across regions, Kozlov *et al.*’s analysis suggests that if ecologists want to use herbaria for this purpose, collectors should be polled to ensure that collector biases do not vary systematically between locations.

Finally, we suggest that Kozlov *et al.*’s conclusion that herbarium specimens may serve as “distorting mirrors” is unjustified because it reaches beyond the limited quantifications of bias in their study. Like any historical reconstruction from fossils^15^ or modern specimens^11^, it is important to identify potential sources of bias in the data. However, a bias or source of noise along any one axes does not preclude the value of these data for exploring variation across other axes. Instead of explaining how the biases they detected across species and curators might affect herbarium-based data analyses, the authors instead appear suggest that these data are too inaccurate to be of use. Kozlov *et al.* do not provide any additional evidence to support this statement. We here draw parallels to the use of paleontological or archeological data. If the detection of taphonomic biases in fossil collections had halted the fields of archeology and paleontology, the vast knowledge we have gleaned from fossil data would never have been collected. We suggest that ecologists approach biases in modern collections data in the same way through the robust assessments of particular biases that are most likely to affect conclusions, rather than a wholesale rejection of such data.

## Conclusion

While biases in herbarium data are well recognized^16,17^, there remains tremendous value in these historical specimens^3,18^. Our re-analysis of Kozlov *et al.*’s data reveals that herbarium specimens provide a useful record of historical herbivory, contrary to the authors published conclusions. We fully support efforts to validate herbivory estimates derived from herbarium samples with gold standard ecological sampling but urge the use of power analyses and/or rarefaction curves to ensure samples sizes sufficient to yield reliable estimates. We also note that while largescale efforts to better quantify collector bias in specimen data across species are useful, we are often more interested in temporal and spatial trends within species to capture the fingerprint of global change (e.g.^19^). We hope that Kozlov *et al.*’s disproportionate and at times unjustified criticism of herbarium specimens as a source of species interactions data will not discourage ecologists from leveraging the information captured within these specimens, which are increasingly being mobilized online and represent a rich resource for inferring past ecologies^20^.

## Author contributions

E.K.M. and T.J.D. contributed to statistical analyses. E.K.M., T.J.D., and C.C.D. wrote the paper.

## Additional information

The authors declare no competing interests.

## Appendix S1: A discussion of least-squared vs. arithmetic means

In Fig. 1, and where appropriate in statistical analyses, we report results from both arithmetic means (Fig. 1a) and the marginal (least-squared) means reported in Kozlov *et al.*’s manuscript (Fig. 1b). The model structure for the calculation of the least-squared means was not described in the original manuscript, and in subsequent conversations, the authors were not able to explain the reason for including these least-squared means (rather than arithmetic means) to us beyond that they account for “uneven sample sizes”. Least-squared means do not necessarily correct for uneven sample sizes, but they can help account for uneven sampling across a particular variable. Given no information, we initially assumed Kozlov and colleagues collected herbivory measurements from multiple sites, and that perhaps these means may have been derived to correct for the uneven application of sampling methods across these sites. However, the vast majority of their species (14/17) were only collected at one site.

We suggest our weighted regression model on the arithmetic means provides the best and most transparent representation of the data. Nonetheless, we provide analyses based on the authors’ reported least-squared means alongside the arithmetic means derived from their raw data. In taking this approach, we aimed to be as inclusive as possible of the authors preferences for their least-squared means estimates, which we struggled to replicate.

---

**Figure S1.**
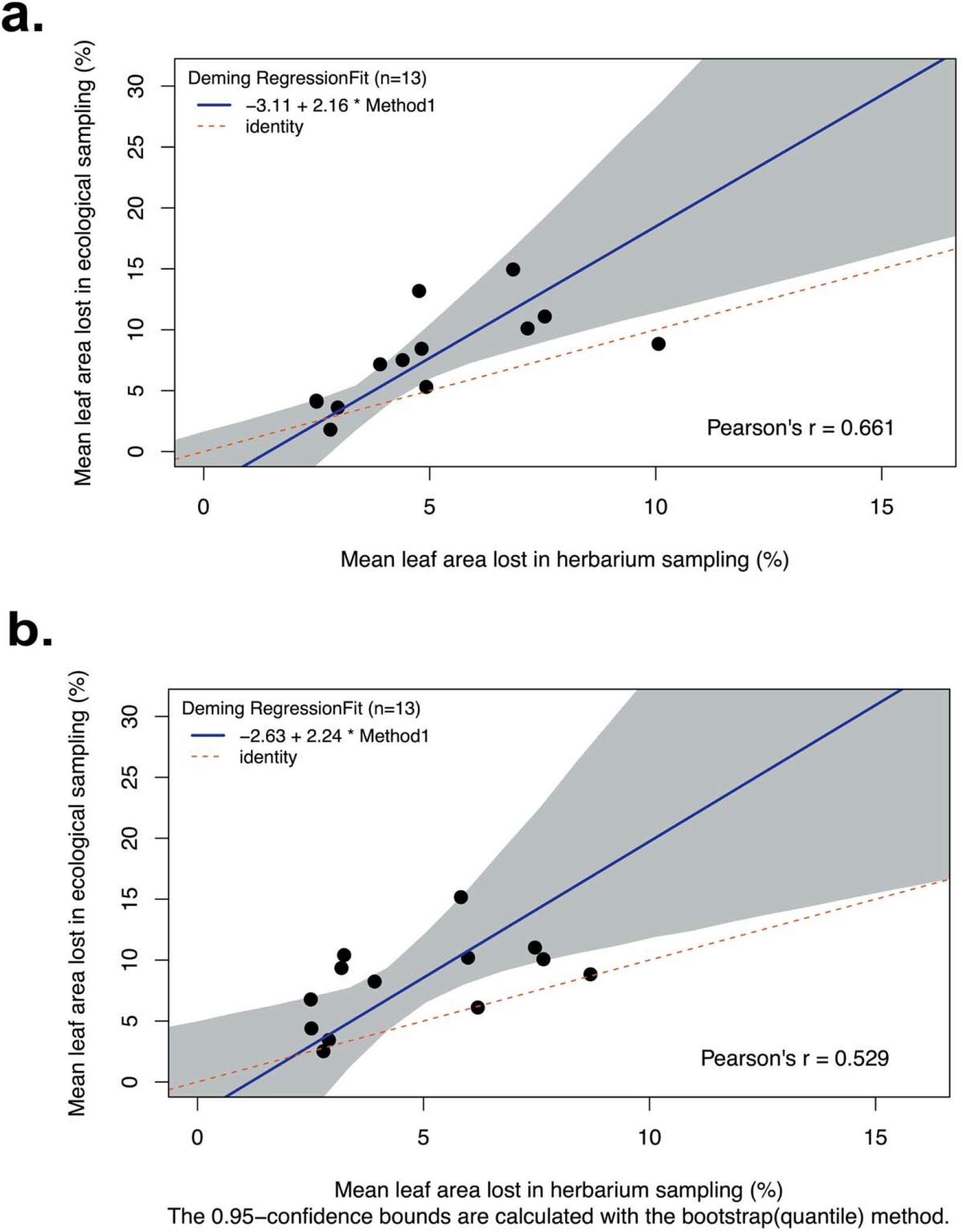
Deming regression fits described in Table 1, Models 7-8.

## Notes

### Competing Interest Statement

The authors have declared no competing interest.

